# A conserved class of viral RNA structures regulate translation reinitiation through dynamic ribosome interactions

**DOI:** 10.1101/2023.09.29.560040

**Authors:** Madeline E. Sherlock, Conner J. Langeberg, Katherine E. Segar, Jeffrey S. Kieft

**Affiliations:** Department of Biochemistry and Molecular Genetics, University of Colorado Anschutz Medical Campus, Aurora, CO, 80045, USA; New York Structural Biology Center, New York, NY, 10027, USA; Innovative Genomics Institute, University of California, Berkeley, CA, USA; Department of Biochemistry and Molecular Biophysics, Columbia University, New York, NY, USA

**Keywords:** viral RNA, reinitiation, ribosome, RNA structure, cryoEM

## Abstract

Certain viral RNAs encode proteins downstream of the main protein coding region, expressed through “termination-reinitiation” events, dependent on RNA structure. RNA elements located upstream of the first stop codon within these viral mRNAs bind the ribosome, preventing ribosome recycling and inducing reinitiation. We used bioinformatic methods to identify new examples of viral reinitiation-stimulating RNAs and experimentally verified their secondary structure and function. We determined the structure of a representative viral RNA-ribosome complex using cryoEM. 3D classification and variability analyses reveal that the viral RNA structure can sample a range of conformations while remaining tethered to the ribosome, which enabling the ribosome to find a reinitiation start site within a limited range of mRNA sequence. Evaluating the conserved features and constraints of this entire RNA class in the context of the cryoEM reconstruction provides insight into mechanisms enabling reinitiation, a translation regulation strategy employed by many other viral and eukaryotic systems.

## INTRODUCTION

Most eukaryotic mRNAs are monocistronic, containing one main open reading frame (ORF) and therefore encoding one functional protein. The first phase in translation of these mRNAs is initiation, which canonically requires numerous eukaryotic initiation factor (eIFs) proteins, and protein synthesis occurs at a start codon near the 5’ end of the mRNA (Aitken and Lorsch, 2012; Hinnebusch and Lorsch, 2012). In the second phase, the ribosome elongates through the coding region until it reaches an in-frame stop codon and terminates, releasing the protein product (Dever and Green, 2012; Hellen, 2018). The ribosome is then recycled for another round of translation by sequential removal of the large (60S) ribosomal subunit, transfer RNA (tRNA), and the small (40S) ribosomal subunit from the mRNA (Young and Guydosh, 2022). However, under specific circumstances this final recycling step can be altered: successful termination and peptide release but incomplete ribosome recycling leads to translation “re-initiation.” In reinitiation, mRNA sequence downstream of the termination codon is translated by the same ribosome as the upstream main ORF, leading to a second protein product expressed from a single mRNA (Gunisova et al., 2018; Sherlock et al., 2023; Skabkin et al., 2013). Reinitiation is an important part of the repertoire of mechanisms that control gene expression at the level of translation but is poorly understood.

Many viruses use reinitiation as a strategy to expand the coding and regulatory potential of a single RNA. Some viral mRNAs contain downstream ORFs that are translated exclusively through programmed translation reinitiation (Ahmadian et al., 2000; Guo et al., 2009; Meyers, 2003; Napthine et al., 2009; Powell, 2010; Powell et al., 2008a; Powell et al., 2008b), and in certain cases, structures embedded within the viral mRNA act in *cis* to promote this (Ahmadian et al., 2000; Luttermann and Meyers, 2007; Powell, 2010; Powell et al., 2011). Importantly, reinitiation is distinct from other RNA structure-based strategies employed by viruses to alter translation, such as internal ribosome entry sites (IRESs), ribosomal frameshifting, and stop codon readthrough (Firth and Brierley, 2012; Firth et al., 2011; Giedroc and Cornish, 2009; Jaafar and Kieft, 2019; Mailliot and Martin, 2018). IRESs can induce translation of a downstream ORF in a bicistronic context, but they do not rely on upstream translation (Filbin and Kieft, 2009; Martinez-Salas et al., 2018; Wilson et al., 2000). With frameshifting and stop codon readthrough the ribosome never terminates, creating fusion proteins with C-terminal extensions (Dinman, 2012; Loughran et al., 2018; Riegger and Caliskan, 2022). In contrast, reinitiation requires that termination proceeds to completion, including peptide release, resulting in two separate functional proteins.

There is currently no standard term to broadly describe reinitiation signals. Therefore, we use “reinitiation-<u>s</u>timulating <u>e</u>lement” (RSE) to refer to sequences and structures capable of inducing programmed reinitiation events. This draws a parallel with the nomenclature for programmed ribosomal frameshifting events, which are induced by frameshift-stimulating elements (FSE).

The importance of RSEs is illustrated by the fact that one specific class was independently identified as the mechanism responsible for downstream expression from bicistronic viral mRNAs in both influenza B virus, a segmented negative-sense single-stranded RNA (-ssRNA) virus in the *Orthomyxoviridae* family, and in members of *Caliciviridae*, a family of positive-sense single-stranded RNA (+ssRNA) viruses (Herbert et al., 1996; Horvath et al., 1990; Meyers, 2003; Powell et al., 2008b). In these viruses, the two ORFs of the bicistronic mRNA slightly overlap and the upstream ORF stop codon and downstream ORF start codon can even share nucleotides (Figure 1A). Despite significant differences in phylogenetic lineage, reinitiation in both of these viral contexts is driven by the same class of RSE, termed termination <u>u</u>pstream ribosome <u>b</u>inding <u>s</u>ite (TURBS) RNA structures (Meyers, 2003).

**Figure 1.**
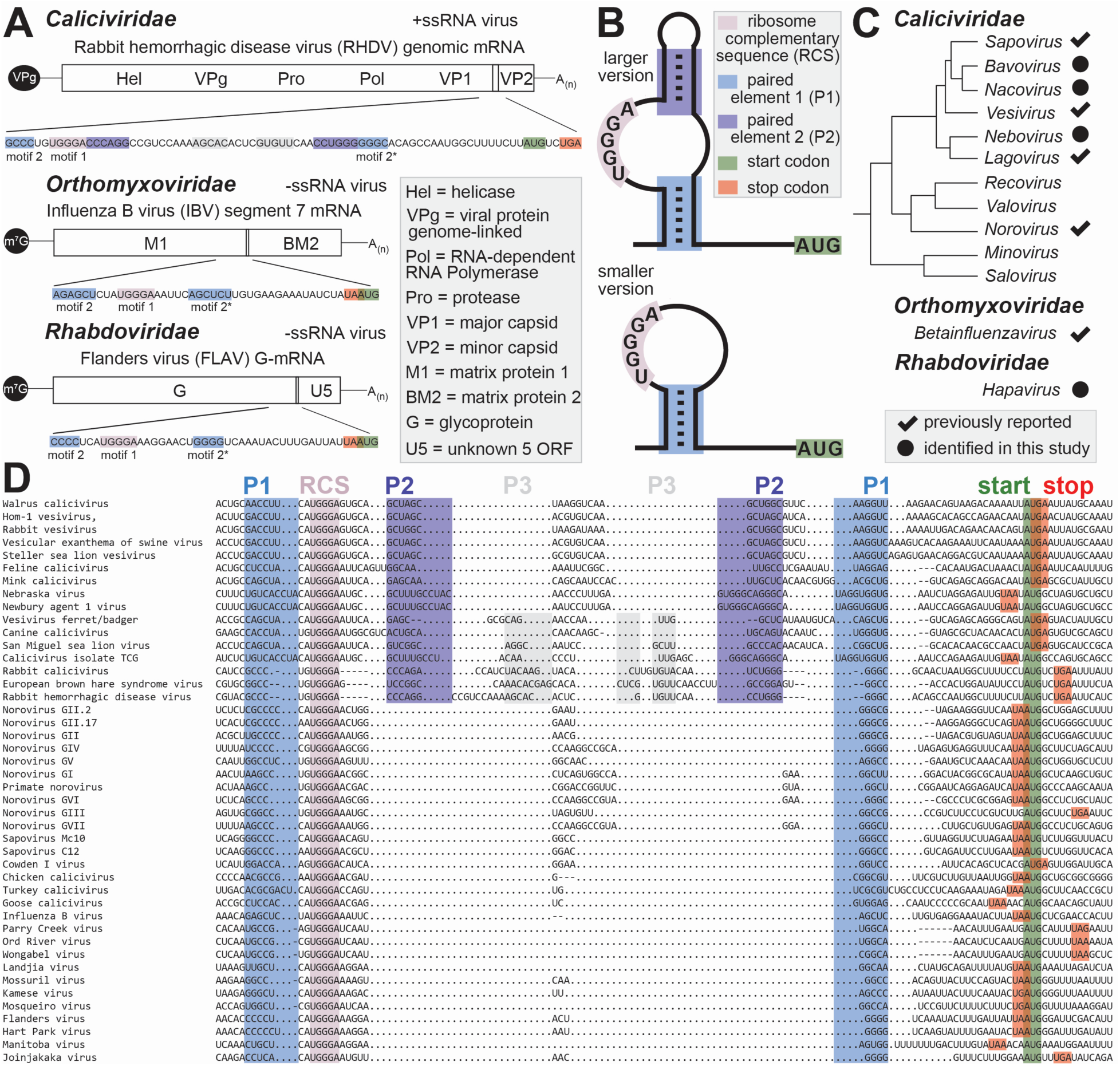
Genomic context and phylogenetic distribution of a class of reinitiation stimulating RNA elements in viruses. (A) Secondary structure cartoons of the two major types of reinitiation stimulating elements, which are located upstream (5’) of the downstream reinitiation start codon. These RNA structures contain either two helical elements (P1 and P2) with the ribosome complementary sequence (RCS) embedded in an asymmetric internal loop (top), or a single helical element (P1) with the RCS located in the apical loop (bottom). (B) Genomic locations of reinitiation stimulating elements in viruses in different families. (C) Phylogenetic distribution of all previously reported and newly identified reinitiation stimulating RNAs in viruses. (D) Alignment of the sequence, secondary structure (paired elements 1-3), and downstream start codon of all RNAs belonging to the termination upstream ribosome binding site (TURBS) class of reinitiation stimulating element RNAs. See also Supplemental File 1 and Table S2.

Within the TURBS class of RSE, mRNA sequences (not the encoded peptide sequence) located 5’ of the stop/start codon overlap region are necessary for reinitiation, while sequences 3’ of the stop/start motif are dispensable (Luttermann and Meyers, 2007; Meyers, 2003; Powell et al., 2008b). Specific portions of the upstream RNA sequence needed for reinitiation are termed motifs 1 and 2 (Luttermann and Meyers, 2007). Motif 1 comprises a stretch of conserved nucleotides complementary to a portion of 18S ribosomal RNA (rRNA) in the 40S subunit, specifically the solvent accessible apex of helix 26, also referred to as expansion segment 7 (ES7). This proposed binding location is well supported by successful induction of reinitiation in yeast only with a TURBS RSE mutated in motif 1 to complement the yeast ribosome’s ES7 apical loop (Luttermann and Meyers, 2009; Powell et al., 2011). Motif 2 comprises two stretches of nucleotides that flank motif 1, with the 5’ portion termed motif 2 and the 3’ portion termed motif 2* (Figure 1A,B). The primary sequence is not conserved, but motifs 2 and 2* are complementary and mutations that break this putative intramolecular base pairing are deleterious to reinitiation (Luttermann and Meyers, 2009). Interestingly, compensatory mutations that restore this base pairing do not fully restore function (Powell et al., 2011; Wennesz et al., 2019), implying that this secondary structure model does not fully explain the nature of this class of RSE RNAs. A 3D structural model for TURBS RSE RNAs accounting for potential tertiary interactions has not been proposed or experimentally determined.

In addition to necessary secondary structural features, the TURBS RSE RNAs have a conserved pattern in their position. Specifically, the structure is always fully upstream of the ORF1 stop codon and ORF2 start codon, but the number of intervening nucleotides varies between different viruses (Powell, 2010). While there is some flexibility in the positioning of the RSE, there are boundaries on the distance between the RSE structure and stop/start codons and on the position of the termination site relative to the reinitiation site (Luttermann and Meyers, 2007). This raises questions as to how the new start codon is found after termination and the involvement of translation factors.

Compared to canonical initiation, TURBS RSE dependent reinitiation requires fewer eIFs (Zinoviev et al., 2015). While mutations of the reinitiation AUG start codon to a near cognate are tolerated, methionine is still incorporated at the N-terminus of the resultant protein, implicating a role for initiator methionyl-tRNA (Met-tRNA_i_) in this type of reinitiation (Luttermann and Meyers, 2007; Napthine et al., 2009). In an *in vitro* reconstituted system Met-tRNA_i_ could be delivered by eIF2, the factor that performs this role in canonical translation initiation (Zinoviev et al., 2015). However, Met-tRNA_i_ can also be delivered by eIF2D or its heterodimeric homolog consisting of multiple copies in T-cell lymphoma-1 (MCT1) and density regulated protein (DENR), all of which play significant roles in ribosome recycling (Young et al., 2018; Young et al., 2021). In the same system, eIF3 is not essential (Zinoviev et al., 2015), contrary to previous claims of its involvement through direct binding (Pöyry et al., 2007). However, supplementary eIF3 does improve reinitiation efficiency, especially in the context of certain mutations or when other factors are absent (Powell et al., 2011; Pöyry et al., 2007; Zinoviev et al., 2015).

Reinitiation is a widespread phenomenon that is clearly an important mechanism of translation control, and the TURBS RNAs are unique compared to both cap-dependent initiation, other types of reinitiation, and other RNA structure dependent strategies of altering translation (Gunisova et al., 2018; Sherlock et al., 2023). This, and the fact that these elements are used by many viruses, motivated further investigation into this mechanism of translation regulation. Previous reports include thorough mutational analyses but were limited to a few representative members of the TURBS RNA class, raising the question of how widespread and diverse TURBS elements are. Other fundamental remaining questions include how these structured RSEs display their recognition sequence to bind to the ribosome, and how they remain tethered to the 40S subunit after termination while simultaneously allowing a new start codon to be found either upstream or downstream of the stop codon.

Here, we combine computational, biochemical, and structural biology approaches to investigate the TURBS class of RSE. We identified several new instances of this class of RSE with diverse reinitiation efficiencies and evaluated the conservation and constraints common to all examples of this class. We then explored the nature of the RSE RNA interactions with the 40S subunit by determining the structure of one representative member in complex with the ribosome using cryo-electron microscopy (cryoEM). Our results provide significant new insight into previously observed characteristics of the TURBS RNAs and contextualizes shared features across the class. In addition, the observation of dynamic motions of the TURBS RSE on the ribosome and molecular modeling allow us to propose a model for how the RSE remains bound to the ribosome while a new start codon is located.

## RESULTS

### Bioinformatic analyses identify new putative viral RSEs

We first sought to understand the distribution and diversity the TURBS class of RSEs by using comparative sequence analysis and homology searches. We aligned sequences from previously studied RSEs from human norovirus (HuNoV), human sapovirus (HuSaV), feline calicivirus (FCV), rabbit hemorrhagic disease virus (RHDV), and influenza B virus (IBV). These represent four genera within the *Caliciviridae* family (*norovirus*, *sapovirus*, *vesivirus*, *lagovirus*) and the *betainfluenzavirus* genus of *Orthomyxoviridae*. The features initially aligned were the ‘UGGGA’ comprising motif 1, hereafter referred to as the ribosome complementary sequence (RCS), the annotated downstream AUG start codon, and base pairing comprising motif 2/2*, hereafter referred to as paired element 1 (P1). For the RNAs proposed to form the larger version, base pairing of a second paired element (P2) was included (Figure 1B). Using this alignment as a seed file, we used the program Infernal (Nawrocki and Eddy, 2013; Nawrocki et al., 2009) to query a viral sequence database, identifying sequences that might represent new TURBS RSE RNAs. Some of these examples were expected, since TURBS elements were proposed to exist in all viruses within the *norovirus*, *sapovirus*, *vesivirus* and *lagovirus* genera of *Caliciviridae* beyond the representative sequence in the initial alignment (Powell, 2010). Other putative TURBS are new examples not previously described (Figure 1C-D, Supplemental File 1, Table S2).

New representatives in the *Caliciviridae* family include Newbury 1 agent virus (N1V), the sole member of the *nebovirus* genus, which is predicted to contain P2 and be a larger version of the structure. Smaller versions of the motif were identified in chicken calicivirus (ChCV), the sole member of *bavovirus*, as well as turkey calicivirus (TuCV) and goose calicivirus of the genus *nacovirus*. All putative new TURBs identified in the *Caliciviridae* family match previously reported genomic context, with the AUG nucleotides in the alignment matching the annotated start codon for the VP2 minor capsid protein. Interestingly, reinitiation appears to be used by most viruses in the *Caliciviridae* family but is not ubiquitous. Viruses in the *recovirus*, *valovirus*, *minovirus*, and *salovirus* genera do not appear to contain an RSE RNA or at least not one consistent with the TURBS class. Notably, viruses in recovirus and valovirus contain the proper stop/start context for reinitiation (UAAUG) but not the matching RNA secondary structural features; these examples are discussed in more detail in the following sections.

Outside of the *Caliciviridae* family there were examples identified in many but not all members the *hapavirus* genus within the *Rhabdoviridae* family, which are non-segmented - ssRNA viruses. Based on the structural alignment by homology searches, all examples in *hapavirus* represent the smaller stem-loop version of the RSE RNA structure. For most of the hapavirus representatives, the genomic context of the putative RSE RNA is in the G mRNA segment, situated near the 3’ end of the first ORF, just upstream of the downstream, out-of-frame U5 gene (Figure 1A). Interestingly, the U5 protein encodes a viroporin (Walker et al., 2011), which is the same class of protein as the BM2 protein of IBV also expressed by reinitiation. In a few of these genomes, the RSE motif lies within the bicistronic N mRNA segment, poised to reinitiate and express the U4 protein, which encodes a small acidic protein of unknown function (Walker et al., 2011).

### Functional assays confirm activity of new putative RSE RNAs from diverse viruses

All putative new examples returned by our homology searches are predicted to be authentic active TURBS RSEs, based on the fully conserved RCS, the presence of P1, and their genetic context. To experimentally validate these new putative sequences as authentic RSE RNAs within the TURBS class, we used a dual luciferase reporter system. This system is designed to quantitatively recapitulate reinitiation events in lysate by testing whether new putative RSE sequences can induce downstream translation on a bicistronic RNA. The reporters contain WT or mutated RSE sequences from a variety of viruses inserted between the upstream *Renilla* luciferase (Rluc) and downstream Firefly luciferase (Fluc) ORFs. The various viral sequences were cloned into the pSGDLuc v3.0 vector, originally developed to accurately measure frameshifting events, which contains StopGo 2A peptide sequences from foot-and-mouth disease virus before and after the insert region (Loughran et al., 2017). This liberates both luciferase proteins from the inserted viral sequence at the protein level (Rluc at its C-terminus; Fluc at its N-terminus) to ensure luminescence is not distorted by added sequence (Loughran et al., 2017). The negative control RNA has a stop codon following the Rluc ORF followed by the Firefly ORF which is in a different frame and lacks a start codon within 170 nucleotides of the Rluc stop codon. The positive control RNA has a small insertion that disrupts the Rluc stop codon, leading to uninterrupted translation of the in-frame Fluc ORF (Figure 2A).

**Figure 2.**
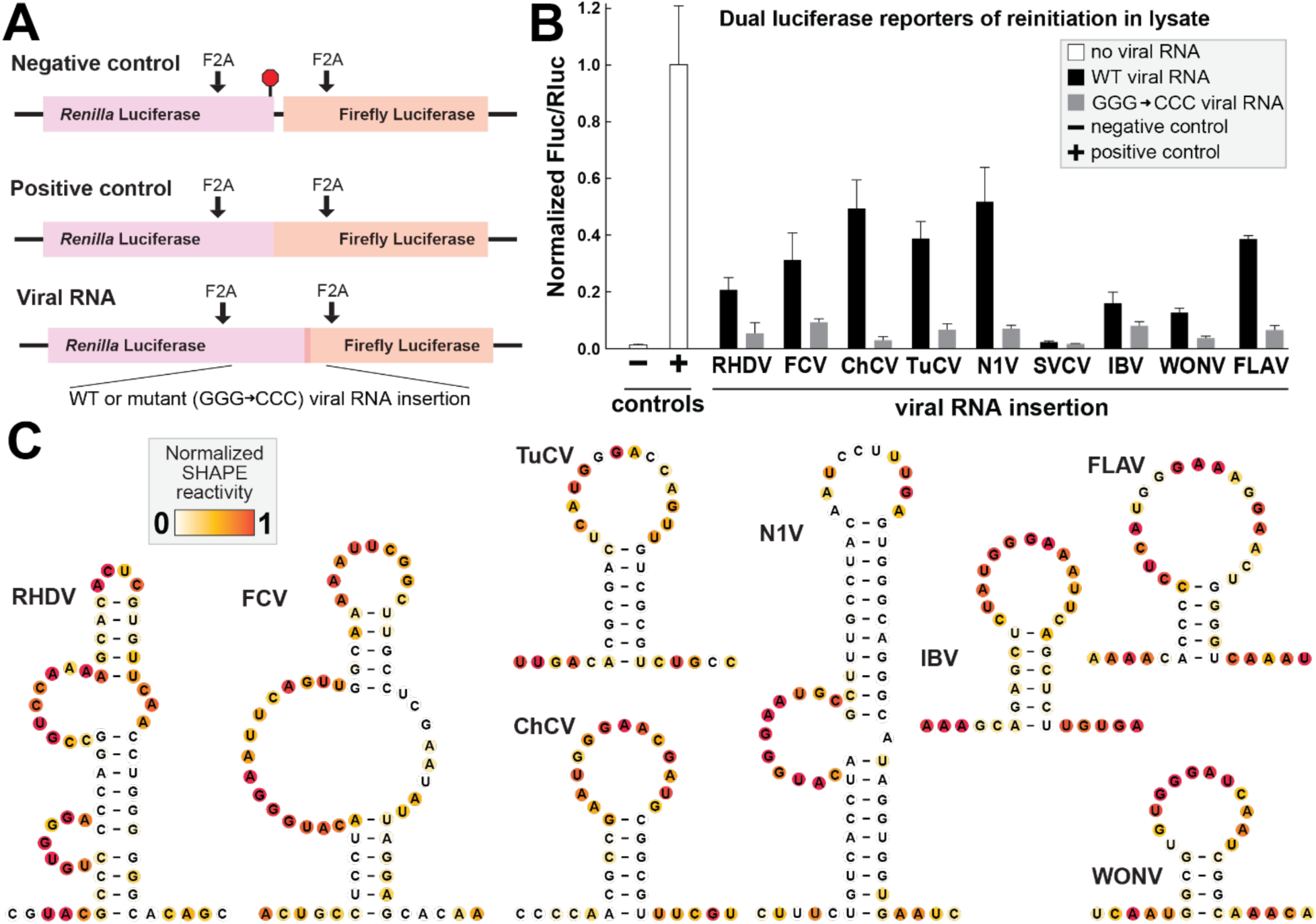
Structure and function of reinitiation-stimulating RNA representatives from diverse viruses. (A) Design of reinitiation reporter constructs, which encode *Renilla* luciferase in the upstream ORF and Firefly luciferase in the downstream ORF. The negative control reporter RNA is derived from the pSGDLuc v3.0 plasmid and contains an in-frame stop codon after the Rluc gene and no in-frame start codon for the downstream, out-of-frame Fluc. The positive control reporter RNA does not contain an in-frame stop codon between the Rluc ORF and in-frame downstream Fluc ORF. Reporters carrying putative viral reinitiation stimulating RNAs were constructed such that the annotated upstream stop codon and downstream start codon were in frame with the *Renilla* and Firefly ORFs, respectively. Mutant viral sequences all contain a GGG → CCC mutation in the RCS predicted by bioinformatic alignment, as depicted in Figure 1D. (B) Translation assays performed in rabbit reticulocyte lysate containing dual luciferase reporters carrying no viral RNA (controls), or putative WT or mutant reinitiation stimulating sequences from rabbit hemmorhagic disease virus (RHDV), feline calicivirus (FCV), chicken calicivirus (ChCV), turkey calicivirus (TuCV), Newbury agent 1 virus (N1V), Saint Valerien calicivirus (SVV), influenza B virus (IBV), flanders virus (FLAV), or wongabel virus (WONV). The ratio of luminescence from the downstream (Fluc) to upstream (Rluc) proteins was normalized to the positive readthrough control. (C) Reactivity to the chemical probing reagent NMIA, assayed *in vitro* by selective 2’ hydroxyl acylation analyzed by primer extension (SHAPE) of putative reinitiation stimulating RNAs mapped onto the predicted secondary structure model. The full sequence that was probed, including flanking hairpins added to normalize reactivity, can be found in Table S1.

Viral sequences inserted into this vector included at least 30 nucleotides upstream of the RSE RNA element (i.e. the 5’ end of the P1 stem), at least three codons of native viral downstream coding sequence, and were designed such that the upstream and downstream luciferase proteins were in the same relative reading frames as in the native viral genomic context. For each putative viral RSE, we tested both the WT and a sequence carrying a mutation to the RCS, specifically a GGG → CCC mutation. Full length dual luciferase RNAs were transcribed *in vitro* and added to lysate to measure translation via the luminescence activity of the two luciferase proteins. The ratio of downstream (Fluc) to upstream (Rluc) luminescence for the positive control RNA was set to 1, as every ribosome that translates the first ORF for this construct should continue through the second ORF. The relative luminescence ratios for all other constructs were normalized to this (Figure 2B). The negative control RNA showed extremely low levels of Fluc luminescence, near background (0.015 ± 0.002).

Using this reporter system, we first validated the previously reported RSEs from RHDV, FCV, and IBV. All resulted in signal corresponding to downstream translation well above the level of the negative control (Figure 2B). Consistent with previous reports of 3-20% efficiencies for these RSEs, the ratio of downstream to upstream translation was lower than the positive control (Luttermann and Meyers, 2014; Meyers, 2003; Powell et al., 2008b). The GGG → CCC mutation, which should prevent binding to ES7 of the rRNA (Luttermann and Meyers, 2009; Meyers, 2003; Powell et al., 2008b), resulted in a decrease in downstream translation, indicating the dual luciferase reporters and *in vitro* assay faithfully report reinitiation events induced by the RNA structure.

We next tested several new putative RSE that we identified through our homology searches, selecting one representative of each new genus in *Caliciviridae* and both genomic contexts in the *hapavirus* genus of *Rhabdoviridae*. For each, the WT sequence induces downstream translation consistent with reinitiation. Likewise, in each case the RCS mutation leads to less downstream translation compared to WT. These results indicate that this translation is driven specifically by the RSE. We note that the level of relative downstream translation comparing mutant constructs across viruses is low but variable. This could be due to some background level of reinitiation, which would likely vary between different stop/start contexts, or spurious internal initiation could contribute since the experiments were performs in a lysate system and not in cells.

As mentioned in the previous section, some viruses within *Caliciviridae* contain a stop/start context at the VP1/VP2 junction that could indicate a reinitiation site, but the homology search did not identify an RSE matching the TURBS class. Still, we tested the sequence corresponding to the region upstream and including the stop/start overlap of St-Valérien virus (*valovirus* genus of *Caliciviridae*) for reinitiation activity. Downstream translation for this WT viral sequence was very low – only slightly above the negative control, confirming this virus indeed does not contain an RSE of this class. Although this virus contains a ‘UGGGA’ sequence >100 nucleotides upstream of the stop codon, a GGG → CCC mutation does not impact the already very low downstream translation level, as expected. This result supports the conclusion that other tested examples are true RSEs, as not all viral sequences drive downstream translation. The expression strategy for VP2 in St-Valérien virus is currently unclear.

### Newly identified RNA structures fold into similar secondary structures

Although the homology searches strongly suggested a common secondary structure used by these newly identified and diverse RSEs, we next confirmed this experimentally. We used *in vitro* structure probing with the chemical modifier N-methylisatoic anhydride (NMIA) to assess the relative flexibility of each nucleotide position within and flanking the RSE structure. We probed examples from RHDV, FCV, and N1V to represent the larger TURBS RSE version with P2 present, and examples from ChCV (*nacovirus*), TuCV (*bavovirus*), IBV (*betainfluenzavirus*), flanders virus (FLAV; *hapavirus*), and wongabel virus (WONV; *hapavirus*) to represent the smaller version of the structure. The probing revealed that all RNAs folded as expected with low reactivity for nucleotides predicted to be involved in the base pairing of P1 and P2, when present (Figure 2C). In all cases, the loops have overall higher reactivity, including the nucleotides complementary to ES7 of the 18S rRNA. This is consistent with these complementary sequences being unpaired and available to bind the ribosome. The RSE from RHDV was previously proposed to contain a third paired element (P3) and this is also by the chemical reactivity profile. Overall, these chemical probing data support the previously proposed secondary structure models used to create the alignments for homology searches and confirm the folding pattern of newly identified examples belonging to this class of viral RSE RNA.

### Conservation and variation among reinitiation-stimulating sequences

After verifying the function and secondary structures of the new TURBS RSEs identified through our bioinformatic searches, we evaluated the conservation and variability among all members. We took the output sequence alignment and created consensus sequence and secondary structure models of both the larger versions and smaller versions (Figure 3A). The P1 and P2 stems are supported by covariation, indicating that the specific nucleotide identities are largely unimportant if base pairing is maintained, with one exception. When aligned from the bottom of P1, the fourth pair is conserved as C-G in all examples, indicating that this position might play a more specific structural role (see Discussion). The P3 stem in RHDV only appears to be present in a few sequences, mostly other members of *lagovirus*, and this helix does not display significant covariation due to this limited distribution.

**Figure 3.**
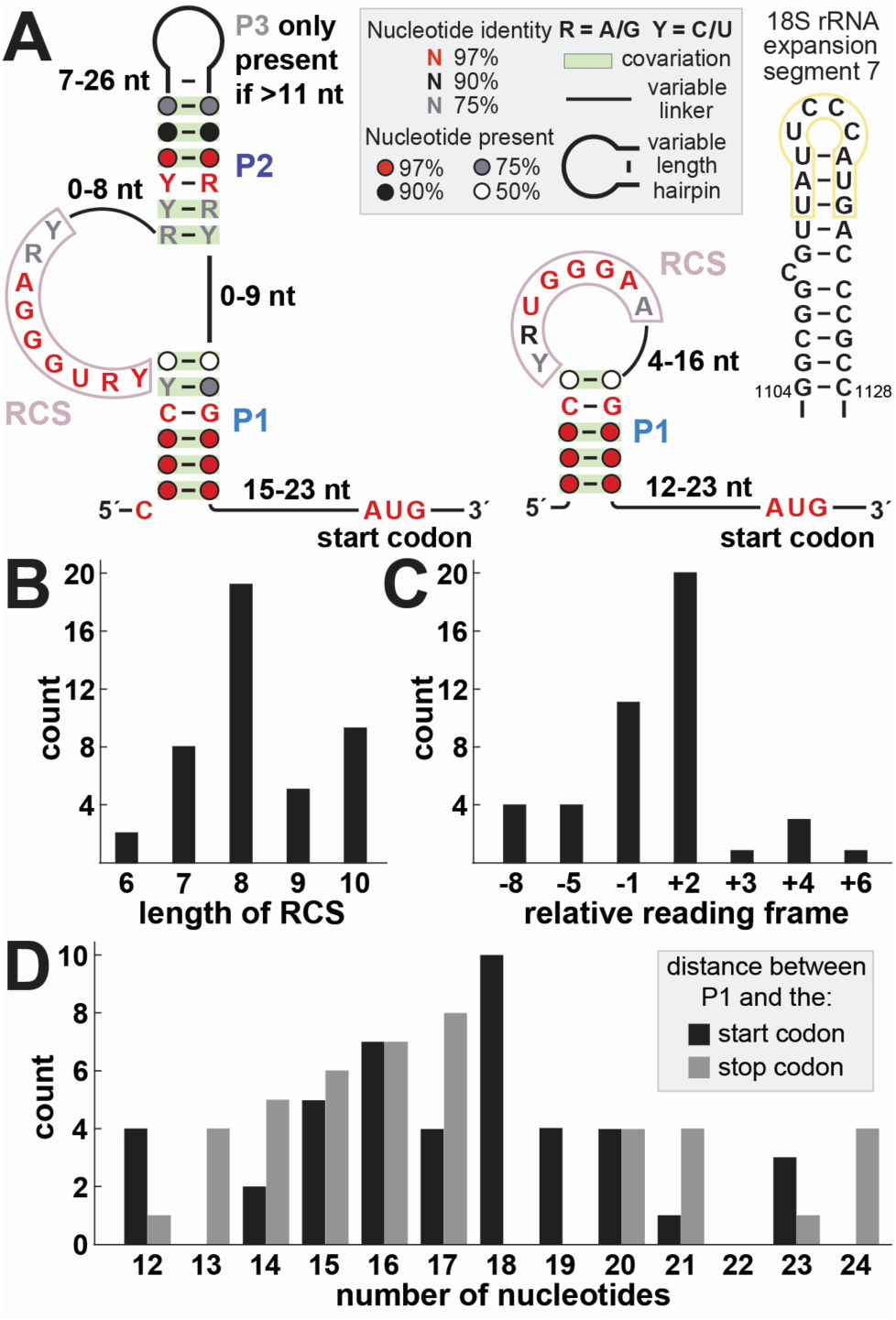
Conserved features among all viral TURBS-like RSE RNAs. (A) Consensus sequence and secondary structure models of all large (left) and small (right) versions of the TURBS class of reinitiation stimulating elements derived from 43 unique viruses. The sequence and secondary structure of the 18S ribosomal RNA from G1104-C1126, a portion of helix 26 also known as expansion segment 7, is depicted with the nucleotides complementary to the RCS within the viral RNA outlined in yellow. See also Supplemental File 1 and Table S2. (B) Histogram of the number of base pairs that could form between the viral RNA and rRNA, as determined by the length of consecutive complementary nucleotides, allowing for G•U wobble pairs. (C) Histogram of all possible reinitiation reading frames, comparing the position of the downstream start codon to the upstream stop codon. (D) Histogram of all possible lengths of the linker between the last nucleotide within the P1 stem and the first nucleotide in either the start (black) or stop (gray) codon among all examples of TURBS RNAs.

Due to the way the alignment was initially constructed, the reinitiation start codon is fully conserved as an AUG and connected to P1 by a linker 12-23 nucleotides in length with no primary sequence conservation. The stop codon of the upstream ORF is always present near the start codon, as discussed in the following section. In the larger examples of this structure, the nucleotide directly 5’ of the P1 stem is absolutely conserved as a cytidine, whereas this nucleotide can have any identity in the smaller examples of the structure. The largest area of primary sequence conservation is the RCS (formerly referred to as motif 1). This element is conserved among all examples with ‘UGGGA,’ but the amount of base pairing possible between the viral RNA and rRNA ES7 spans a larger region in some examples. When allowing for G•U wobble pairs, the RCS is always at least six nucleotides in length as nucleotide preceding the UGGGA is always a purine that can pair with U1120 of the rRNA. At most, the complementary region comprises ten nucleotides (Figure 3B), potentially pairing to positions 1112-1121 of the 18S rRNA (Figure 3A, highlighted in yellow).

### Some reinitiation sequences have extended base-pairing with rRNA

The observation that some TURBS potentially make more base pairs with rRNA was made previously (Luttermann and Meyers, 2009; Powell, 2010), but it was not experimentally demonstrated. To test, we performed chemical probing of the RSE RNAs from IBV, RHDV, and FCV in the presence of an *in vitro* transcribed RNA corresponding to nucleotides 1104-1120 of the 18S rRNA, representing the apical portion of ES7. We then subtracted these reactivities from those measured for the viral RNA alone to create a differential reactivity profile (Figure 2C), revealing which nucleotide positions become more or less flexible in the presence of the rRNA stem loop (Figure S1). For the IBV RSE, which has the potential to form eight base pairs with ES7, only seven nucleotides display a decrease in reactivity in the presence of ES7 RNA. The RHDV RSE could form seven base pairs with ES7, six of which exhibit decreased reactivity, while the FCV RSE could form ten base pairs with ES7, nine of which exhibit decreased reactivity. Therefore, the number of nucleotides in the RCS might not always reflect the number of nucleotides that base pair with the rRNA, although some pairs could form and be more dynamic. The ability to form up to nine or ten base pairs in some cases is interesting because ES7 contains an apical tetraloop of only 4 nucleotides. Therefore, to form more than 4 intermolecular base pairs with the viral RNA, part of the ES7 stem must unwind (Figure S1). Furthermore, the proposed intermolecular base pairing requires that at least one nucleotide for the RSE RNA and at least one nucleotide of ES7 rRNA are unpaired, presumably to form linkers to relieve geometric constraints imposed by the other structured elements.

### Constraints on RNA structure, termination site, and reinitiation site positioning provide insights into mechanism

To gain additional insight into the mechanism of reinitiation by TURBS RSEs, we now examined their genomic context to include the location of the structured element compared to the stop codon and start codon, as well as the relative locations of these two codons. For each example, we identified the annotated stop codon, ensuring that this was the first in-frame termination codon for the upstream ORF, and recorded its position within the alignment relative to the reinitiation start codon (Figure 1D). The most common stop/start codon contexts were <u>UA**A**</u>**UG**, <u>UG**A**</u>**UG**, **A<u>UG</u>**<u>A</u>, and <u>UGA</u>NN**AUG**, where N represents any nucleotide identity, the stop codon is underlined, and the start codon is in bold (Table S2). All three canonical stop codons are found in this class of RNAs, indicating any type of termination event is compatible with reinitiation.

We used the distance between the upstream ORF termination site and the downstream reinitiation start site to calculate the relative reading frame for each example, setting the stop codon as the zero position (e.g. <u>UA**A**</u>**UG** is in the +2 frame, **A<u>UG</u>**<u>A</u> is in the -1 frame). This revealed a range of allowable relative reading frames from -8 to +6 for RSEs in this class (Figure 3C). The numerical values for the relative reading frames in the context of reinitiation differ from that of ribosomal frameshifting due to the relationship between the termination and reinitiation sites. For reinitiation, the relative reading frames are offset by three nucleotides in terms of how far the ribosome must traverse relative to the mRNA. Upon termination, the stop codon occupies the A-site of the decoding groove whereas the reinitiation start codon must be placed in the P-site to pair with Met-tRNA_i_. Therefore, reinitiation in the +2 frame (e.g. <u>UA**A**</u>**UG**) actually requires the mRNA to move by five nucleotides within the decoding groove of the ribosome for the start codon to be properly placed. In contrast, reinitiation in the -1 frame (e.g. **A<u>UG</u>**<u>A)</u> only requires movement by 2 nucleotides. This likely explains why the reinitiation reading frames skew toward the negative, with nearly all examples accessing reading frames between -8 and +4.

Certain reinitiation frames are fundamentally incompatible with the identities of canonical start and stop codons. For example, the -2 frame cannot be used in reinitiation because the third nucleotide of the start codon must be a guanosine (AUG) while the first nucleotide of the stop codon must be a uridine (UAA/UGA/UAG). Codons that are incompatible due to nucleotide identities are -2, 0, and +1, while the -3, -6, and -9 relative reading frames would lead to reinitiation events that immediately terminate due to being in frame with the upstream ORF stop codon. While the upstream and downstream ORFs cannot be in the same reading frame if the reinitiation start site is upstream of the termination site, reinitiation in the same frame is allowable if the reinitiation start codon is fully downstream of the termination site, such as +3 or +6. Indeed, both arrangements are represented in the sequence dataset in both members of the *nacovirus* genus.

A histogram of the number of nucleotides between the closing of P1 and the start or stop codon shows that the range and distribution of linker lengths are similar for both codons, either 12-23 or 12-24 nucleotides for the start and stop codon, respectively (Figure 3D, Table S2). The lower limit is likely imposed by the footprint of the ribosome, being the minimum distance spanned from the decoding groove to the outside of the mRNA exit channel where the RSE interacts. The upper limit on the linker length is likely imposed by a need for the RNA structure to be near its ES7 binding site when the stop codon is in the A site. The importance of this is supported by previous experimental data showing reinitiation cannot be achieved if the start codon is too far downstream of the RSE RNA structure (Luttermann and Meyers, 2007). There are likely complex kinetic and thermodynamics at play enforcing this restriction on the upper distance limit, especially in the context of structural data presented in the following sections.

### Binding site and architecture of a TURBS RSE RNA structure revealed by cryoEM

Our bioinformatic and biochemical analyses yield a more complete picture of TURBS RSE diversity and conservation, but raise several other questions, including: How does an RSE of this class align itself on the 40S subunit and stay tethered while allowing a new start codon to be found? How do constraints on codon position relate to TURBS function? To answer questions regarding molecular basis of RNA structure-induced reinitiation, we determined the structure of a TURBS RSE bound to the ribosome.

We incubated purified 80S ribosomes with *in vitro* transcribed RHDV RNA (6951-7039) to assemble a complex that was then used for single particle cryoelectron microscopy (cryoEM). Following preliminary analysis and classification (Figure S2), extra map features were readily identified on the 40S subunit in the ES7 region, which we tentatively assigned to the RHDV RNA (Figure 4A). Most of the good 80S particles had this added feature, suggesting most ribosomes were bound to RHDV RNA. Subtracting the 60S signal then masking for the 40S subunit revealed higher resolution details of the site of interaction (Figure 4B). The extra map feature on ES7 extends well beyond what is accounted for by the apo ribosome structure (Figure 4C) with two lobes extending away from the 40S surface. Although the overall resolution of the complex was ∼3Å, the local resolution of the map representing the TURBS RSE was considerably lower, similar to an rRNA expansion segment that extends away from the 40S surface.

**Figure 4.**
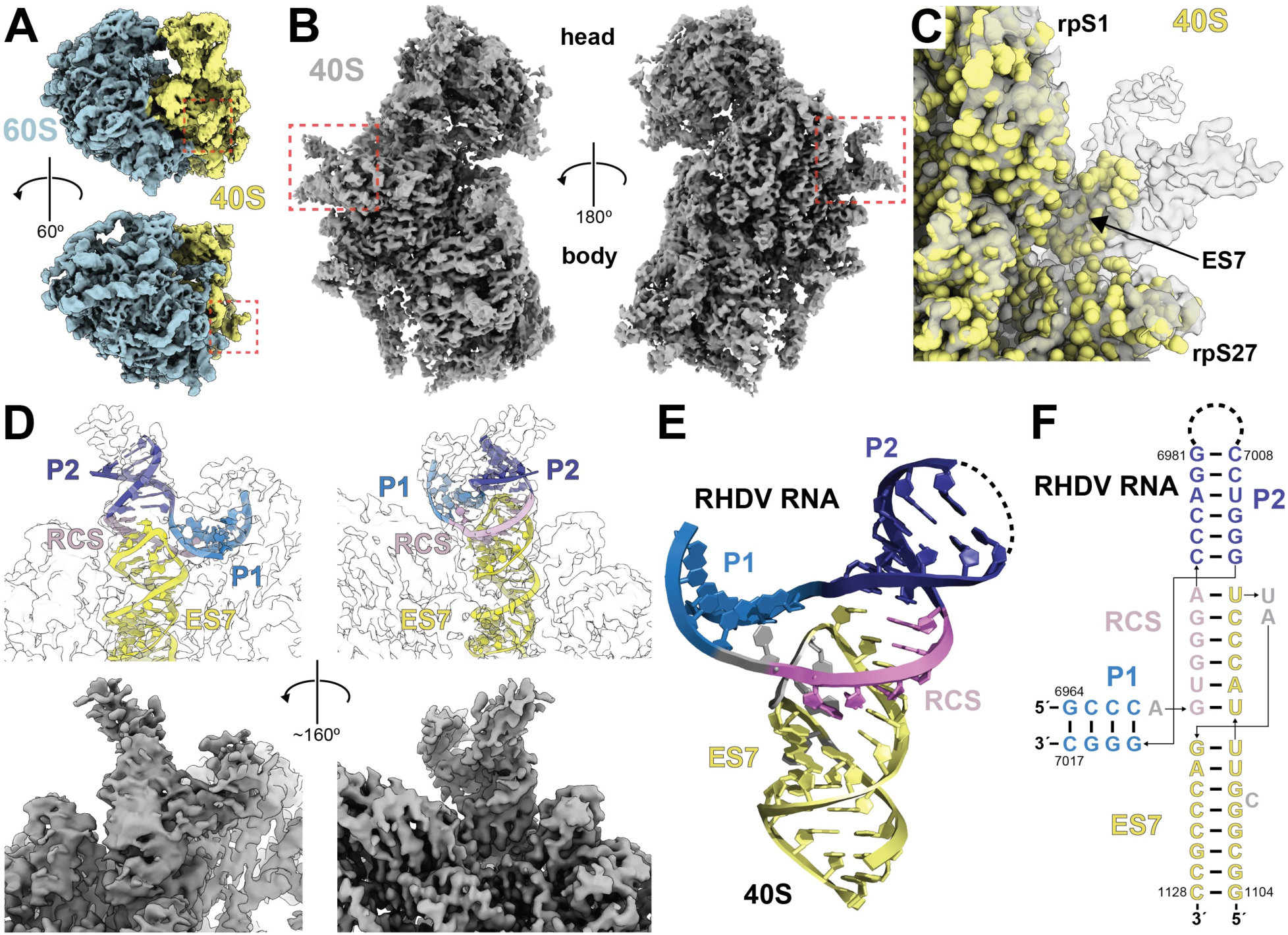
CryoEM reconstruction of a RSE-ribosome complex reveals the binding site and general architecture of the RNA structure. (A) Refined map of the full RHDV RNA-80S complex prior to masking. Density corresponding to RHDV RNA is boxed in red. (B) Refined map of the masked 40S subunit of the RHDV RNA-ribosome complex. Density corresponding to RHDV RNA is boxed in red. (C) CryoEM map of the RHDV RNA-ribosome complex, zoomed in on the region boxed in red in panel B. An apo-40S structure (yellow, PDB: 6ZMW) was fit into the map, revealing extra density corresponding to the RHDV RNA contiguous with ES7. (D) A model of the RHDV RNA (top) built into the density of a locally refined map (bottom), including intermolecular base pairs between the ribosome complementary sequence (RCS) of RHDV and the apical loop of ES7. (E) Model of the RHDV RNA with ribosomal ES7. Nucleotides in gray (RHDV A6969; 18S rRNA U1119, A1120, C1109) could not be unambiguously built into the density. (F) Secondary structure of the RHDV RNA bound to the apical portion of ES7, as determined by the model in panel E.

### Structural model of the viral RNA in complex with a mammalian ribosome

Guided by the chemical probing data, secondary structure model, and proposed binding scheme between the RHDV and ES7 (Figures 2C, S1), we determined the overall architecture of the RNA structure and assigned P1 and P2 each to one lobe of the map that extending away from the ribosome surface. Once the general topology of the RHDV RNA was clear, we built a model of the rRNA-RSE RNA complex into a locally refined map (Figure 4D-F). We fit an existing 40S structure to place the base of ES7, for which the map matched very closely to previous models. The distal portion of ES7 (nucleotides G1110 to C1123) diverged from existing structures, starting with minor variations then propagating to major differences further away from the core of the ribosome; these residues were built manually into the density (depicted in yellow in Figures 4-6). The helix of ES7 is disrupted by a bulged nucleotide, C1109, for which the map is somewhat ambiguous and the local resolution is poor (Figure 5A), and this nucleotide was modeled according to a previous structure (Brito Querido et al., 2020).

**Figure 5.**
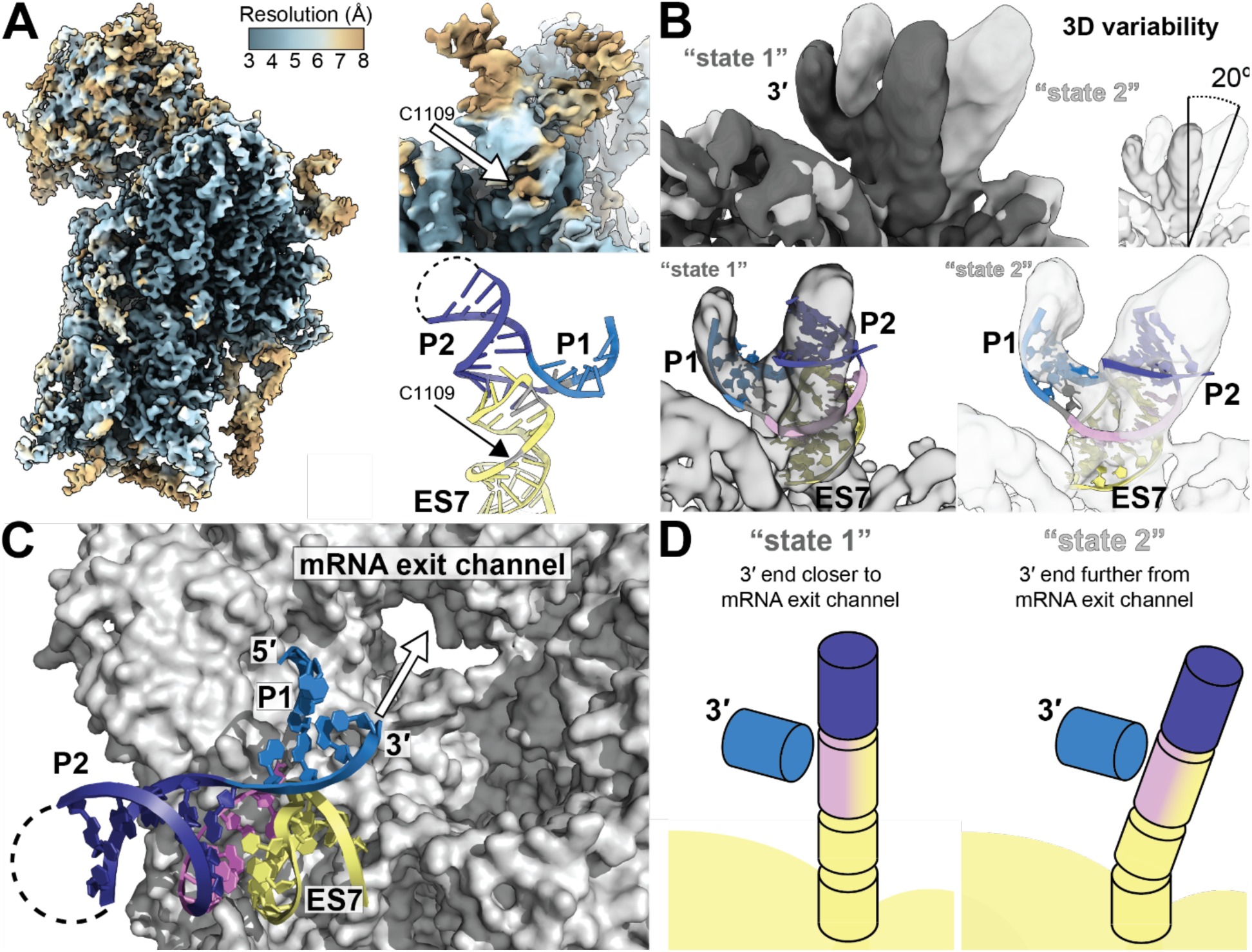
Conformational heterogeneity reveals dynamics related to reinitiation start site selection. (A) Local resolution of the masked, refined 40S-RHDV RNA map. The position of 18S rRNA C1109 is depicted within the local resolution map and the model with the same view. (B) Maps representing the two extreme states of 3D variability among the particles, which differ in their orientation by 20°. The model of RHDV RNA bound to the apical portion of ES7 rRNA (G1110 to C1123) was fit into the two extreme states. (C) Surface view of the 40S with the model of RHDV RNA bound to ES7, with the view looking down into the mRNA exit channel from the decoding groove. (D) Simplified model of the orientation of helices in the two extreme states of 3D variability.

**Figure 6.**
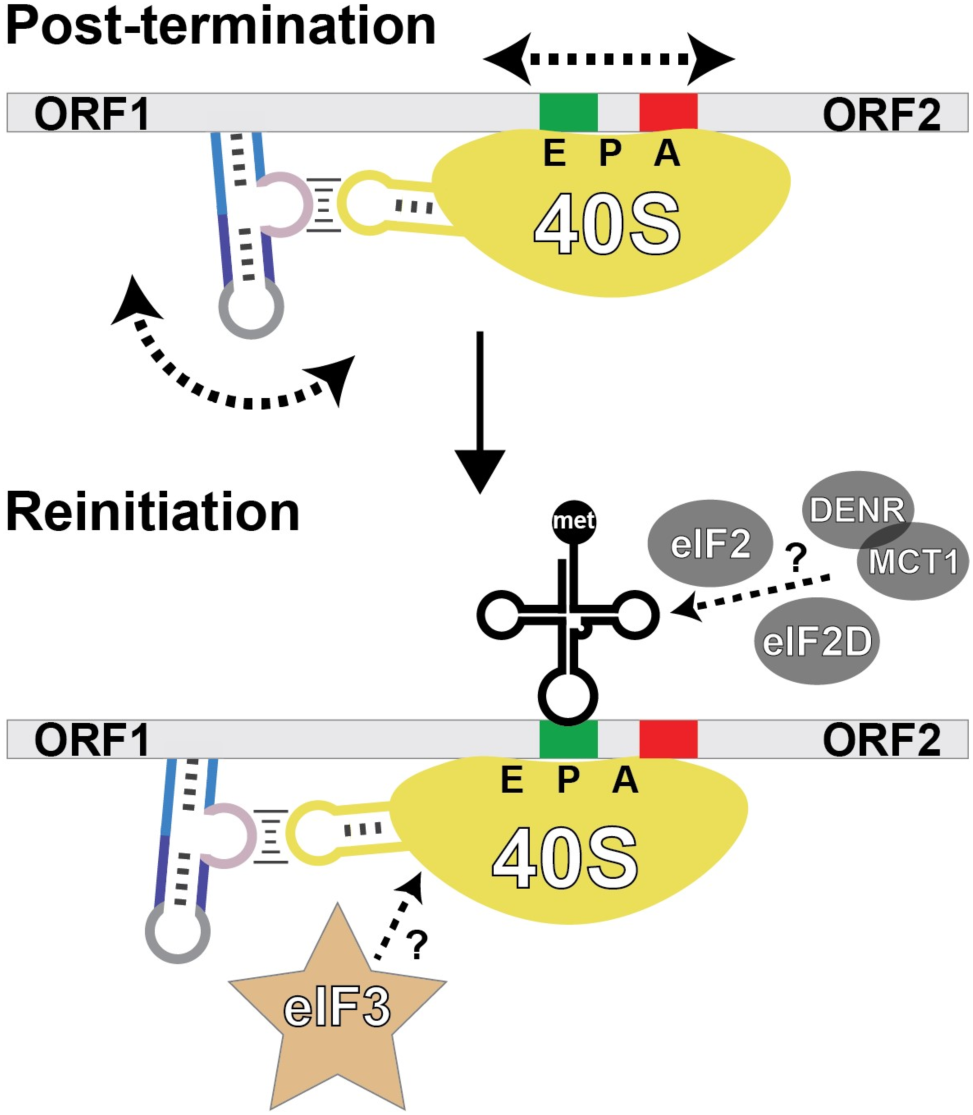
Model for the structural bases for reinitiation start site selection within a limited window. During termination, the stop codon is in the A-site. We propose that conformational flexibility and dynamics allow the RHDV RNA to remain bound to ES7 of the 40S while sampling different states. This motion moves the 3’ end of the structure, which could propagate to the downstream mRNA in the decoding groove and allow sampling of nearby RNA including the nearby reinitiation start codon in the P-site. Other factors, including eIF3, could select for certain conformations of RHDV-ES7 and influence start site selection. Initiator tRNA^Met^ is used in this type of reinitiation which can be delivered by eIF2, eIF2D, or DENR/MCT1 *in vitro* (Zinoviev et al., 2015), but the mechanism(s) of delivery *in vivo* have not yet been defined.

The intramolecular base pairing within ES7 was maintained through the U1112-G1121 pair, at which point the helix continues with little disruption to the A-form geometry but consists of intermolecular base pairs between the rRNA and RHDV RNA (depicted in lilac in Figures 4-6). The location of contiguous density from intermolecular base pairs to the lobe corresponding to P1 along with chemical probing data support the pairing of six nucleotides between the internal loop of RHDV and ES7, leaving A6969 unpaired (Figure 4E,F). This requires unwinding of the two top base pairs normally formed in ES7, with U1113 and A1114 reconfiguring from intra- to intermolecular base pairing while U1119 and A1120 become unpaired (Figure 4F). The continuous stacking from ES7 into the RSE leads to a structure in which the RSE appears as an “extension” of the ES7, pointing away from the ribosome surface.

Although the nature of the ES7-RHDV RNA base pairing is clear, the local resolution of the map in other regions, especially the two lobes extending away from the ribosome surface, was insufficient to unambiguously build all portions of the structure (Figure 5A). The location and orientation of the P1 and P2 helices within the two lobes is clear due to the strand directionality necessary to achieve the RHDV RNA-rRNA base pairing. However, within the lobes some features such as the distinctive phosphodiester backbone, or major and minor groove were not clear. Therefore, we built P1 and P2 by placing A-form helices at an angle best matching the map and refined their location and connectivity to satisfy proper backbone geometry. The precise location of the two unpaired nucleotides of ES7 rRNA (U1119, A1120) and A6969 of the RHDV RNA are also ambiguous given the resolution in those regions. These nucleotides (depicted in gray) were thus modeled consistent with the density and satisfying favorable backbone angles. The unpaired nature of these nucleotides and lower local resolution suggest these are flexible regions of the RNA (see Discussion). This is consistent with chemical probing data, as A6969 of RHDV is reactive to in both the unbound and bound forms (Figure 2C, S1).

Beyond the features described above, the maps did not allow us to unambiguously build other parts of the RHDV RNA without overspeculation. Specifically, the nucleotides that form the P3 stem and a large internal loop were not visible, likely due to substantial conformational heterogeneity; this is supported by high chemical reactivity of the loop nucleotides (Figure 2C). Although some additional map features are present in the area adjacent to P1 beyond the eight nucleotides comprising the helix, it is poorly resolved (>8 Å) and it is unclear which nucleotides might occupy this space based solely on chemical probing data and the consensus model. As such, did not model any nucleotides near the 5’ end of P1 (G6964). Finally, none of the nucleotides upstream or downstream of P1 were built into the model, again due to this ambiguity.

### The orientation of the RHDV RSE on the ribosome aligns with the decoding groove

The binding site of the RHDV RSE on ES7 places it directly adjacent to the part of the decoding groove where mRNA exits after being translated. Presumably, the RNA comprising the RSE passes through the ribosome, then folds and interacts with ES7 as termination is occurring on a stop codon that is in the decoding groove in the A site. The structure of the bound RHDV RSE is consistent with this, as the P1 stem is positioned such that it is aligned well with the decoding groove. Specifically, the 3’ end of this stem points at the decoding groove where the downstream RNA is placed during a termination and reinitiation event. Notably, there are no map features present in the decoding groove that would correspond to the RNA between the 3’ end of P1 and the stop/start codons, although this sequence was present in the RNA used to make the complex. We speculate that this could be due to the way in which the sample was generated using purified 80S ribosome and RNA only. Nonetheless, the position and orientation of the RHDV RSE on the 40S explains how it binds and is connected to the stop and start codons in the decoding groove during termination and reinitiation, respectively.

### Dynamics in the RHDV RSE-ribosome complex move the viral RNA relative to the decoding groove

The position and orientation of the RHDV RSE RNA on the ribosome show how it is linked to RNA in the decoding groove, and analysis of the dynamics reveal how it could stay “tethered” to the ribosome while allowing a new start codon to be found. Specifically, using 3D variability and 3D classification, we found that the viral RNA can occupy a range of positions while remaining bound to the rRNA (Figure 5B). These positions do not appear to be fully discrete states, as increasing the number of 3D classes leads to more states populated along the axis of this inferred motion. This motion means that the small number of particles classified into each minor state significantly limits the resolution, resulting in increasingly poor map quality in the P1 and P2 stems compared to the RCS-ES7 helix (Figure 5A). We speculate that the nucleotide positions that enable these dynamics are the unpaired, flexible residues which have low local resolution and whose positions could not be definitively placed. In particular, the bulged C1109 within the rRNA appears to be poised to affect the trajectory of the helix by introducing a kink in the backbone (Figure 5A). Other nucleotides that are more peripheral but likely also enable dynamics are A6969 of RHDV, which would specifically allow flexibility in the position of P1, and the unpaired linker of ES7 (U1119, A1120).

Examining the nature of the dynamic motions more closely, it appears the extended helical stack of rRNA, then viral RNA-rRNA hybrid, then viral RNA (P2), can maintain all base pairs but sample different local conformations, which propagate to large differences in the location of peripheral regions of the structure. The two extreme states identified by variability analysis demonstrate a 20° difference in positioning of the RSE RNA relative to the overall ribosome structure (Figure 5B). That is, the RHDV RNA remains bound but “rocks” back and forth on the ribosome’s surface. This leads to a 13 Å difference in the location of the 3’ end of P1 between the two states, but we cannot exclude the possibility that other, even more widely separated states exist but are populated very rarely.

We propose that the conformational heterogeneity observed in the cryoEM data reflect important dynamics for the reinitiation function of this structure, possibly explaining how the RSE RNA stays bound to the ribosome while a start codon is found upstream or downstream of the stop codon. The directionality of the inferred motion changes the distance between the RSE structure and the mRNA exit channel (Figure 5C,D). Specifically, movement of the P1 structure relative to the ribosome could allow movement of the downstream RNA in the decoding groove such that it could reposition relative to the ribosome (Figure 6). The dynamics revealed by the cryoEM data would allow different codons to be sampled in the P-site after termination. We explore the implications of this motion and model in the Discussion.

## DISCUSSION

Translation reinitiation is an important strategy for controlling translation, used by many viruses to expand the coding potential of their genomes and to produce proteins necessary for viral propagation in defined ratios. Here, we explored the structure and function of a class of structured viral RNA sequences that promote reinitiation. Using a combination of comparative sequence analysis and homology searches, biochemistry, and structural biology, we determined the sequence diversity and structural conservation of these RNAs and explored their interactions with the ribosome. Our structure reveals how the RNA can stay bound to the ribosome surface while undergoing conformational dynamics that likely are important for reinitiation. We propose a model for how this class of RNA can facilitate reinitiation that is consistent with previously published data and points the way toward the next set of questions and experiments.

### Proposed identity and interactions for ambiguous regions of the cryoEM reconstruction

In all cryoEM maps, there were features in the map adjacent to P1 that could not be accounted for by the nucleotides comprising that stem. While we did not build into these features, we speculate that the flanking nucleotides on the 5’ end of P1 are likely candidates for occupying this space. Among larger TURBS RSE RNAs, the nucleotide immediately preceding P1 (C6963 of RHDV) is always a cytidine (Figure 3A). Systematic truncations of the upstream sequences for both IBV and RHDV indicate that multiple nucleotides 5’ of the P1 stem are necessary to achieve full downstream translation efficiency (Meyers, 2003; Powell et al., 2008b). For IBV, the two nucleotides prior to P1 display low chemical probing reactivity both in the ES7-bound and unbound forms (Figure 2C, S1), but mutation of the nucleotide immediately upstream of P1 (C727) to an adenosine causes both 5’ nucleotides to display high reactivity while base pairing of P1 remains intact (Figure S3). These observations suggest that the C immediately upstream of P1 is involved in important tertiary contacts. If this nucleotide makes tertiary contacts with other portions of the RSE, these could contribute to these mysterious map features.

The structural basis for conservation of the fourth base pair of P1 as a C-G pair (Figure 3A) is also currently unexplained. Mutations of the P1 stem of RHDV such that base pairing is maintained but with different nucleotide identities, including the C6967-G7014 pair, causes a >80% reduction in reinitiation activity compared to WT, which was attributed to misfolding into an alternate secondary structure (Wennesz et al., 2019). While misfolding could contribute, we propose that this specific C-G is needed for tertiary contacts, perhaps related to conserved C6967. These interactions might position RSE elements correctly in 3D space or could prevent misfolding. Understanding the nature of the intra- or intermolecular interactions formed by these nucleotides motivates future experimental investigations into their role, including high-resolution structure determination.

### Possible interactions between the RSE and ribosomal proteins

Another set of putative interactions between the RHDV RNA and the 40S subunit that are suggested by the cryoEM analysis are with neighboring ribosomal proteins, which may depend on the position of the RNA. Specifically, there appears to be continuous density between the regions occupied by RHDV RNA and regions occupied by ribosomal proteins. “State 1” (Figure 5B) puts nucleotides of RHDV within the 5’ portion of the RCS near residues 199-203 of ribosomal protein S1 while “state 2” puts nucleotides in the 3’ portion of the RCS and rRNA close to residues 72-75 of ribosomal protein S27. Due to the overall architecture, these proteins would likely interact with the RNA backbone and not rely on specific nucleotide identities in these positions, but the details of these potential interactions are ambiguous due to the local resolution.

### Viral RSEs and IRESs share a binding site, but differ in their mechanism

The TURBS class of viral RSEs is not the only viral RNA structure known to interact with the ES7 portion of rRNA to manipulate translation. The hepatitis C virus (HCV)-like class of IRESs contains a stem loop which also base pairs with the loop of ES7. Specifically, C1116-C1118 of the ES7 loop base pair with three consecutive guanosines in the loop of HCV IRES domain IIId, while U1115 interacts with a bulged nucleotide in a different portion of IRES domain III (Kieft et al., 2001; Quade et al., 2015). Notably, these IRES-rRNA interactions are only with nucleotides in the loop of ES7 and no major structural rearrangements within ES7 are necessary for binding. In contrast, our work shows that binding of the RHDV RSE to rRNA requires unwinding of two base pairs, leading to substantial disruptions to the structure of the ES7 stem loop compared to the apo ribosome or IRES-ribosome complex (Figure S4). These disruptions may also allow the ES7 stem to be more dynamic when bound to a TURBS RSE versus the HCV IRES.

Aside from their shared binding location on the ribosome and ability to induce a translation event independent of certain eIFs, there are significant differences between TURBS RSEs and HCV-like IRESs in both structure and function. The HCV-like class of IRESs comprises a much larger and more complex structure (Asnani et al., 2015; Kieft et al., 1999), enabling interactions with other regions of the ribosome. In particular, domain II of the HCV IRES reaches into the decoding groove, interacting with the E site to help position the start codon in the P site to achieve internal initiation (Brown et al., 2022; Filbin and Kieft, 2011; Quade et al., 2015). It is thus unsurprising that cryoEM reconstructions of the HCV IRES bound to the ribosome show density within the decoding groove (Brown et al., 2022). In contrast, our maps of the RHDV RSE bound to the ribosome do not show any density in the mRNA channel. While this could be due to construct design or technical limitations, the lack of density could reflect an important mechanistic distinction. That is, reinitiation requires a ribosome first translates the upstream ORF, since the RSE is unable to “de novo” place mRNA in the decoding groove on its own (Horvath et al., 1990; Luttermann and Meyers, 2007; Napthine et al., 2009). Therefore, the RSE RNA structure can only initiate translation after termination (but before full ribosome recycling) when the mRNA is already properly positioned.

Previously, it was proposed that the RSEs do not function as IRESs in part because the ribosome must translate through the upstream structure and allow it to refold before proper RSE RNA-rRNA interactions can form (Napthine et al., 2009; Powell et al., 2008b). While the kinetics of co-translational refolding could certainly impact the efficiency or frequency of reinitiation, we argue that this is likely not the major factor in distinguishing between reinitiation by a TURBS RSEs and HCV-like IRESs. One argument against this post-translation refolding model is that in both chemical probing experiments and in our cryoEM maps, the viral RNA (including sequences flanking the structured element) can bind the ribosome without a specific type of refolding event. We favor the model of RSEs lacking the ability to de novo place mRNA in the decoding groove as a distinguishing characteristic compared to an IRES.

### eIF3 binding is compatible with a viral RSE-ribosome complex

While eIF3 is not necessary to achieve reinitiation *in vitro*, it increases the reinitiation efficiency and leads to an increase in fidelity of authentic AUG start codon selection in concert with eIF1 (Zinoviev et al., 2015). Furthermore, components of the eIF3 complex were found to crosslink with a RSE RNA (Pöyry et al., 2007). To gain insight into potential roles for eIF3 in reinitiation, we modeled an existing structure of eIF3 bound to the 40S subunit (Brito Querido et al., 2020) onto our map and model of the RHDV RSE-80S complex (Figure S5A). The bound RHDV RNA does not inhibit any interactions between eIF3 and ribosomal proteins, as previously proposed (Zinoviev et al., 2015).

In our model, the subunits closest to the RSE structure are eIF3a and eIF3c, which bind ribosomal proteins S1 and S27, respectively, but do not interact directly with ES7 (Brito Querido et al., 2020). The lobe corresponding to P2 is nestled against the bundle of alpha helices of eIF3c (Figure S5A), and while they are in proximity, there do not appear to be steric clashes. The lobe corresponding to P1 clashes with some of the alpha helices corresponding to the N-terminal portion of eIF3a. While these appear to sterically hinder binding of eIF3 when the RSE is present, there are factors that could alleviate these clashes. Most of the map that overlaps eIF3a is unaccounted for by P1, which as we speculated above, may be occupied by flanking 5’ nucleotides – these might interact with eIF3. The steric clashes could also be overcome by structural rearrangements of eIF3a, the RSE RNA structure, or both. If eIF3 can bind the ribosome while the RSE RNA is present, this represents another difference compared to HCV-like IRESs, which directly bind eIF3 (Kieft et al., 2001; Sizova et al., 1998) but reposition many of its subunits (including eIF3a and eIF3c) away from their canonical binding site on the 40S (Hashem et al., 2013).

### Could eIF3 binding influence reinitiation start site selection?

The position of bound eIF3 in relation to bound RSE RNA and the motion of RSE on the ribosome suggest a mechanism by which the factor could facilitate proper placement of the new start codon after termination. Specifically, the regions of the RSE map that clash with eIF3 are different when comparing the position of the two extreme states revealed by 3D variability analysis (Figure S5B). This observation raises the intriguing possibility that binding of eIF3 post-termination stabilizes a specific conformation of the RSE. By selecting a specific position of the dynamic RSE on the ribosome, binding of eIF3 could position P1 such that downstream mRNA in the decoding groove is in the proper location to allow the start codon to be identified and enable reinitiation. This model predicts the presence of direct molecular contacts between the RSE RNA and eIF3.

This proposed conformational fit model for eIF3’s role in this type of reinitiation contextualizes previously reported data related to this factor and might alleviate seemingly conflicting reports. It was found that eIF3 is pulled down by the RSE RNA from FCV in a manner consistent with an HCV-like IRES, which was proposed to be caused by a direct binding event (Pöyry et al., 2007). However, in this same context a construct carrying a mutation of the RCS is unable to pull down eIF3, strongly suggesting that 40S subunit binding is necessary for crosslinking, in contrast to the proposed direct binding model (Pöyry et al., 2007). If direct interactions do form between the RSE RNA and eIF3, it would likely be in other regions of the viral RNA and the affinity of the viral RNA for eIF3 might be too weak to measure via crosslinking unless both are bound to the 40S subunit. In a reconstituted system with purified factors *in vitro*, eIF3 was not necessary to assemble an initiation complex at the reinitiation start site, but its addition increases the level of reinitiation (Zinoviev et al., 2015). These data are consistent with our proposed role in stabilizing a reinitiation-competent conformation and influencing start codon placement. Ultimately, additional experiments or structures will be necessary to elucidate molecular interactions, binding, and any influence on RSE RNA position by eIF3.

While this structure provides some insight into mechanism and the role of eIF3, it does not inform on the roles of other translation machinery in inducing reinitiation. While Met-tRNA_i_ is a necessary component for this type of reinitiation, it can be successfully delivered by multiple factors (Zinoviev et al., 2015), and the influence of different conformational states of the RSE RNA on its arrival are unclear (Figure 6).

### Model for finding a reinitiation start site within a limited window

We hypothesize that the conformational dynamics of the ribosome-bound RHDV RNA are a shared feature among this entire class of viral RSEs. However, the extent of the conformational heterogeneity and space sampled by different TURBS RSE RNAs might differ. This, in turn, could enable different distances and directions of mRNA movement within the decoding groove to place the reinitiation start codon. Differential dynamics among members of this class might be caused by several factors, including variable lengths of flexible linkers connecting paired elements (Figure 3A), different numbers of base pairs between the RCS and ES7 rRNA, and the absence of the P2 stem. These same factors likely play into the overall measured reinitiation efficiency, which differs between viruses (Figure 2B) (Luttermann and Meyers, 2007; Meyers, 2003; Powell et al., 2008b). In addition, the thermodynamic stability of the RSE RNA structure, its folding kinetics, and the kinetics of termination at different stop codon identities likely play into these observed differences. Future studies that query a more diverse set structures should enable firm conclusions as to the relative contributions of each of these numerous factors.

The distribution of distances between of P1 and the start codon being subject to the same constraints as that between P1 and the stop codon suggests that mRNA in the decoding groove can move relative to the ribosome within a limited range of nucleotides. The upper limit on the start codon location implies that the reinitiating complex does not use 5’ to 3’ scanning, consistent with previous observations (Zinoviev et al., 2015). The overall range of distances between P1 and the start or stop codon of 12-24 nucleotides (Figure 3D) likely reflects the footprint of the ribosome, on the lower end, and a maximal distance beyond which the RCS of the nascently translated and refolded RSE RNA can no longer find its binding site on ES7, on the upper end. It is then noteworthy that the allowable relative reading frames have a stricter upper limit of nine nucleotides (for reinitiation in the +6 frame) with most examples only requiring a shift spanning the distance of five or fewer nucleotides in the decoding groove (-8 to +2 frame) (Figure 3C). The limits on allowable relative reading frames likely reflect the maximal dynamics of the bound RSE structure that enable sampling of different nucleotides in the P-site. Overall, bioinformatic and cryoEM data support our proposed model for reinitiation start site selection within a limited number of nucleotides in either direction enabled by conformational dynamics (Figure 6). This model warrants future exploration to uncover the mechanistic details underlying this type of reinitiation.

## EXPERIMENTAL PROCEDURES

### Bioinformatics

Sequences of five previously reported examples of reinitiation stimulating RNAs in the TURBS class from were retrieved from the National Center for Biotechnology Information (NCBI) sequence database (genome accession numbers: NC_039476.1, NC_002210.1, NC_001481.2, NC_001543.1, NC_010624.1). The sequences were manually aligned in Stockholm format based on the following four features: motif 1 (‘UGGGA’), motif 2 (5’ portion of the P1 base pairing), motif 2* (3’ portion of the P1 base pairing) and the annotated start codon for the downstream ORF (‘AUG’). This seed alignment file was used to query the RefSeq viral database (Brister et al., 2015), downloaded on 01/19/2021, containing one representative sequence for each virus using the program Infernal version 1.1 (Nawrocki and Eddy, 2013), allowing for an E-value cutoff of 1 (instead of the default 0.05) to find potentially enable identifying sequences with more variation. Iterative searches followed by manual adjustments to the alignment were performed until no new sequences were identified. In total, sequences from 44 total unique viruses were returned that match the TURBS class of RSE RNAs (Table S2, Supplemental File 1). The alignment was subsequently separated into two files based on the presence or absence of the P2 stem, which was added manually to the alignment for the larger versions of the structure. The two consensus models were calculated using R-scape (Rivas et al., 2017), visualized with R2R (Weinberg and Breaker, 2011), and adjusted and labeled in Adobe Illustrator (Figure 3A). Various features of the structure (e.g. linker lengths, base pairing potential) were calculated manually and histograms were plotted and labeled in Adobe Illustrator.

### Plasmid construction and RNA preparation of dual luciferase reporter constructs

Dual luciferase constructs containing intervening viral RNAs of interest were made using the pSGDLucV3.0 vector (Addgene 119760). Insertion sequences were ordered as double-stranded DNA gene fragments (IDT) then amplified via PCR. Vector and inserts were prepared by digesting with BglII (NEB) in NEBuffer r3.1 (NEB) at 37 °C for 2 hours followed by digestion with PspXI (NEB) in rCutsmart™ Buffer (NEB) and incubated at 37 °C for an additional 2 hours. The vector was purified via agarose gel purification while inserts were purified directly from the restriction digest reaction, both using the Wizard SV Gel and PCR Clean-up System (Promega). Inserts were incorporated into the vector via ligation reaction consisting of inserts and vector in a 5:1 molar ratio, T4 ligase (NEB), and T4 ligation buffer (NEB). The reaction was incubated at 16 °C overnight then transformed into DH5⍺ *E. coli* and screened for successful ligation via colony PCR. Sequences were verified via whole plasmid sequencing prior to use. the positive control and each viral mutant plasmid was constructed using the Q5 site-directed mutagenesis kit (NEB) according to manufacturer protocol and verified via whole plasmid sequencing. Double-stranded DNAs approximately 5 kb in length, corresponding to both luciferase genes separated by control or viral sequence, were produced via PCR as a template and their size verified by 1.5% agarose gel electrophoresis. These PCR products were then used as templates for *in vitro* transcription with the MEGAscript™ T7 transcription kit (Invitrogen). RNA was purified using the Monarch RNA clean-up kit (NEB) and RNA length and quality was verified via 10% acrylamide gel.

### Translation assays using dual luciferase reporters in lysate

Each reaction contained 500 ng of RNA, which was initially heat re-folded by incubating in 33mM HEPES (pH 7.5) at 95 °C for 1 minute then cooling to room temperature for 5 minutes at which time MgCl_2_ was added to a final concentration of 10 mM. The final 50 µL translation reaction included the 5 µL RNA mixture described above and components from the Rabbit Reticulocyte Lysate system (Promega): 35 µL of rabbit reticulocyte lysate, amino acid mixture minus cysteine (final concentration 16.7 µM), amino acid mixture minus methionine (final concentration 16.7 µM), amino acid mixture minus leucine (final concentration 16.7 µM), and KOAc (final concentration 150 mM). Reactions were incubated at 30 °C for 4 hours. After incubation, reactions were diluted 4X with Passive Lysis Buffer (Promega) and the reaction was split into two technical replicates, each with a final volume of 100 µL. Luciferase activity was measured with a GloMax^®^-Multi Detection system using the Dual-Glo^®^ Luciferase Assay System (Promega). Results were analyzed using Excel (Microsoft) software with measurements from at least three independent experiments, using the average of the two technical replicates from each trial. These measured values were used to calculate the ratio of Firefly (downstream) to *Renilla* (upstream) translation (Fluc/Rluc), then these ratios were normalized with the positive control (in-frame readthrough) set to 1. Data were plotted and labeled using Adobe Illustrator.

### RNA preparation for chemical probing and cryoEM

Template DNA was acquired as gBlock DNA fragments (IDT) and subsequently cloning into the pUC19 vector. Double-stranded DNA (dsDNA) templates, featuring an upstream T7 promoter, were amplified in PCR reactions under the following conditions: 100 ng plasmid DNA, 0.5 µM forward and reverse DNA primers (see Table S1), 500 µM dNTPs, 25 mM TAPS-HCl (pH 9.3), 50 mM KCl, 2 mM MgCl_2_, 1 mM β-mercaptoethanol, and Phusion DNA polymerase (New England BioLabs). Successful amplification of dsDNA was confirmed by 1.5% agarose gel electrophoresis. Subsequently, RNA transcription reactions were carried out using the following conditions: approximately 0.1 µM template DNA, 10 mM NTPs, 75 mM MgCl_2_, 30 mM Tris-HCl (pH 8.0), 10 mM DTT, 0.1% spermidine, 0.1% Triton X-100, and T7 RNA polymerase. These reactions were incubated at 37 °C overnight. Following transcription, insoluble inorganic pyrophosphate was removed through centrifugation at 5000xg for 5 minutes, after which the RNA-containing supernatant was ethanol precipitated using 3 volumes of 100% ethanol at -80 °C for a minimum of 1 hour. The mixture was centrifuged at 21000xg for 30 minutes at 4 °C to pellet the RNA, and the ethanolic fraction was carefully decanted. The RNA was resuspended in 9 M urea loading buffer and then purified via denaturing 10% polyacrylamide gel electrophoresis (PAGE). RNA bands were detected via UV shadowing and subsequently excised. These bands were then extracted by crush soaking in diethylpyrocarbonate-treated (DEPC) milli-Q water at 4 °C overnight. The RNA-containing supernatant was concentrated using spin concentrators (Amicon) to achieve the desired RNA concentration in DEPC-treated water. Finally, the RNAs were stored at -80 °C, with working stocks stored at -20 °C.

### *In vitro* chemical probing of RNAs

RNA structure probing experiments using the SHAPE reagent NMIA were conducted in accordance with established protocols (Cordero et al., 2014). Briefly, 240 µM RNA was refolded by heating to 90 °C for 5 minutes, followed by cooling to ambient temperature and incubation with MgCl_2_ for 20 minutes. The refolded RNA was incubated with NMIA for 15 minutes at ambient temperature, with the following modification conditions: 120 nM RNA, 6 mg/mL NMIA or DMSO, 50 mM HEPES-KOH (pH 8.0), 10 mM MgCl_2_, and 3 nM 6-fluorescein amidite 5’-labeled FAM-RT primer (see Table S1). To quench the modification reaction, NaCl was added to a final concentration of 500 mM, and Na-MES buffer (pH 6.0) was adjusted to 50 mM. Modified RNA was recovered using oligo-dT magnetic beads (Invitrogen Poly(A)Purist MAG Kit). After recovery, the RNAs were washed twice with 70% ethanol and resuspended in water. cDNA was synthesized using SuperScript III (Invitrogen) at 48 °C for 1 hour, following the manufacturer’s instructions. The RNA was subsequently degraded by the addition of NaOH to a final concentration of 200 mM and heating to 90 °C for 5 minutes. Hydrolysis was stopped using an acid-quench solution containing 250 mM NaOAc (pH 5.2), 250 mM HCl, and 500 mM NaCl. DNA was then recovered using magnetic beads, washed twice with 70% ethanol, and eluted in GeneScan 350 ROX Dye Size Standard (ThermoFisher) containing HiDi formamide solution (ThermoFisher). The resulting 5’-FAM-labeled reverse-strand DNA products were analyzed via capillary electrophoresis using an Applied Biosystems 3500 XL instrument. Data analysis to calculate reactivity values at each nucleotide position was performed using HiTrace RiboKit (https://ribokit.github.io/HiTRACE/) (Kim et al., 2013; Kladwang et al., 2014; Lee et al., 2015; Yoon et al., 2011) in MatLab (MathWorks). For differential chemical probing, normalized reactivities values in the presence of ligand were subtracted from normalized reactivities values in the absence of ligand. Figures were generated using RiboPaint (https://ribokit.github.io/RiboPaint/) in MatLab then subsequently labeled in Adobe Illustrator. SHAPE reactivity data was mapped onto the predicted secondary structure model based on the alignment.

### Ribosome purification from lysate

Nuclease-treated bulk rabbit reticulocyte lysate (Green Hectares) was spun down at 20,000 RPM for 15 min to remove debris, nuclei, and mitochondria (Belin et al., 2010). Clarified lysate was filtered with a 0.22 µm filter (Millipore). The supernatant was loaded on to a 30% sucrose cushion (20 mM Tris-HCl pH 7.5, 2 mM MgOAc_2_, 150 mM KCl, 30% w/v sucrose) and ultracentrifuged for 17.5 hours at 36,000 RPM (50.2 Ti rotor) at 4 °C to obtain a ribosomal pellet (Matasova et al., 1991). The pellet was washed and resuspended in a buffer containing 20 mM Tris-HCl pH 7.5, 6 mM MgOAc_2_, 150 mM KCl, 6.8% w/v sucrose, 1 mM DTT, 1 μL RNasin from Promega (Cat N2618). To remove non-resuspended particles, the resuspended pellet was centrifuged again at 10,000 g for 10min at 4 °C and the supernatant was isolated. 15-30% sucrose gradients were prepared using a buffer containing 20 mM Tris-HCl pH 7.5, 2 mM MgOAc_2_, 150 mM KCl, and either 15% or 30% w/v sucrose using the Gradient Master (Biocomp). Gradients were cooled to 4 °C before use. Pellet supernatant was loaded onto gradients and ultracentrifuged at 19,100 RPM for 17.5 hours (SW-28 rotor) at 4 °C. Gradients were fractionated using the Piston Gradient Fractionator and Fractionator Software v8.04 (Biocomp), monitoring for absorption at 260 nm. Fractions corresponding to pure 80S ribosomes were pooled and concentrated to an A_260_ of 95 using an Ultra-15 centrifugal filter unit with a nominal molecular weight limit of 100 kDa (Amicon). This concentrated stock was subsequently diluted to 250 nM using buffer containing 20 mM Tris-HCl pH 7.5, 2 mM MgOAc_2_, 150 mM KCl and 20 μL aliquots were flash frozen in liquid nitrogen and stored at -80 °C.

### cryoEM sample preparation and image acquisition

RHDV RNA at 20 μM was heat denatured at 95 °C and allowed to cool slowly to ambient temperature, then 1 μL of the RNA solution was added to a 20 μL aliquot of 250 nM 80S (described above) and incubated at 37°C for 15 minutes. Using a FEI Vitrobot mark IV, a 3 μl aliquot of the RHDV-80S solution was applied to grids (holey carbon grids C-flat 1.2/1.3 400 mesh VWR) that had undergone plasma cleaning for 6 seconds in a mixture of O_2_ and H_2_ by a Gatan Solarus 950 (Gatan, Inc). Grids were blotted for 3.5-5 seconds at a force of -5, then plunge frozen in liquid ethane.

Movies were collected using a Thermo Scientific 300 kV Krios G3i microscope equipped with a Falcon 4 Direct Detection Camera and Selectris Imaging Filter. A total of 6,589 micrographs were collected (429 at 30° stage tilt to overcome preferred orientations). Each movie consisted of 267 frames collected over 7.76 seconds with a total dose of 49.94 e^-^/ Å^2^. EPU (Thermo Fischer) was used to monitor data collection and specify defocus values in the range of -0.4 to -1.8 μM. Data were collected at 130,000X magnification with a physical pixel size of 0.97 Å and filter slit width of 10 eV.

### cryoEM image processing and map generation

Cryo-EM data were processed using cryoSPARC 4.0 (Punjani et al., 2017). An overview of the processing workflow can be found in Figure S2. Movies were imported then processed (patch motion correction, patch CTF estimation) using default parameters. The 80S portion of an existing structure (PDB: 4UJD; (Yamamoto et al., 2014) was used to generate templates for particle picking. Particles were extracted from micrographs with a box size of 800 x 800 pixels, then downsampled 4X (200 x 200 pixel) to speed up initial classification and refinements. Six initial rounds of 2D classification were performed to remove “junk” particles (e.g. incorrectly-picked particles, ice contamination, aggregates). Particles from all good classes were used for *ab initio* 3D reconstruction requesting two classes followed by heterogeneous refinement resulting in one class with good 80S density and another demonstrating good 60S density but poor 40S density and those particles were removed. Four additional rounds of 2D classification were performed to remove particles belonging to classes with poor resolution and/or alignment.

All resulting good particles were subjected to *ab initio* 3D reconstruction and non-uniform refinement (Punjani et al., 2020), resulting in a map with excellent 60S resolution and relatively poor 40S resolution, as expected due to known dynamics of the small subunit. Nevertheless, density was observed in the 40S in the region of the 40S near the mRNA exit channel beyond what was accounted for by the ribosome and in a similar position compared to IRES structures that bind directly to ES7 (Yamamoto et al., 2014). These particle locations underwent local motion correction (Rubinstein and Brubaker, 2015) then were re-extracted without downsampling. Masks were created for the 60S and 40S subunit, the 60S signal was subtracted, and the masked 40S underwent non-uniform refinement. A mask was generated for the local region surrounding the RHDV RNA (“RHDV mask”) using UCSF Chimera (Pettersen et al., 2004). Focused 3D classification was performed with the RHDV mask using varying numbers of classes. Ultimately six classes resulted in four classes with “good” density in the region corresponding to RHDV RNA and two classes with poor density, which were excluded.

The remaining good 61,643 particles underwent non-uniform refinement, first with the 40S mask then subsequent non-uniform refinement with the RHDV mask. Resolutions for each of these refinements (Figure S2) were estimated using the gold standard Fourier shell correlation (GSFSC) of 0.143. The RHDV mask was also used to perform masked 3D variability analysis (Punjani and Fleet, 2021) for all good particles with a resolution threshold of 8 Å. Initially, multiple modes of variability and resolution thresholds were requested, and one particular mode was always observed, as reported in Figure 5B.

### Model building and refinement

The 40S portion of a previously-solved structure (PDB: 4ZMW; (Brito Querido et al., 2020) was fit into the density of the map resulting from the non-uniform “40S mask” refinement using ChimeraX (Pettersen et al., 2021). All regions of our map agreed with the previously solved model except the 18S rRNA nucleotides G1110-C1123 (i.e. the distal portion of ES7). These nucleotides along with the well-resolved portions of RHDV RNA were built into the density of the “RHDV mask” locally refined map. First, base-paired regions were modeled as A-form helices, starting with 18S rRNA base pairing between 1110-1112 and 1121-1123, then the intramolecular pairing between 18S rRNA 1113-1118 and RHDV 6970-6975, then P1 and P2 of RHDV, which were all sequentially fit into the density using ChimeraX. Backbone connectivity and refinement as well as the placement of nucleotides A6969 of RHDV and 1119-1120 of 18S rRNA were performed using COOT (Emsley and Cowtan, 2004). Modelling of the structure into the two extreme states of 3D variability analysis (Figure 5B) was accomplished by fitting only this portion of our structural model (RHDV RNA, 18S rRNA 1110-1123) into each state using ChimeraX. The distance separating these two states was calculated in ChimeraX by measuring the distance between the N1 of G6964 of RHDV modeled into each state. The angle measured between the two states was calculated by ChimeraX using the position of G6981 (top of P1) in each state with G1121 of the 18S rRNA serving as an anchor point.

Modeling of eIF3 was accomplished by simply adding back all subunits of this complex, which were present in the structure used to build the 40S portion of our structure (PDB: 4ZMW). Modeling of the HCV IRES-ribosome interactions compared to RHSV RSE-ribosome structural model was accomplished by fitting the 40S portion of an existing structure (PDB: 5A2Q; (Quade et al., 2015) into our map then adding back the IRES RNA. Figures of cryoEM maps and structural models were made using ChimeraX, with the exception of Figure 4 panel E and Figure 5 panel C which were created using PyMol (Schrödinger, LLC) and labeled using Adobe Illustrator.

## SUPPLEMENTAL INFORMATION

Supplemental materials (five figures, three tables, one file) are available for this Article.

## AUTHOR CONTRIBUTIONS

MES: Conceptualization, Data curation, Formal Analysis, Funding acquisition, Investigation, Methodology, Visualization, Writing – original draft, Writing – review & editing. CJL: Investigation, Formal Analysis, Writing – review & editing. KES: Investigation, Writing – review & editing. JSK: Funding acquisition, Supervision, Visualization, Writing – review & editing.

## ACKNOWLEDGEMENTS

We thank Salimah Thaxton, Nadine Ramirez, and Andrea MacFadden for technical assistance in cloning and additional members of the Kieft laboratory for helpful discussions. We thank Charles Moe at the University of Colorado Boulder Krios Electron Microscopy Center for assistance with cryoEM data collection. The pSGDlucV3.0 was a gift from John Atkins (Addgene plasmid # 119760). This work was supported by National Institutes of Health (NIH) grant R35GM118070 to JSK. MES was supported by a Jane Coffin Childs Postdoctoral Fellowship. KES was supported by NIH grant T32GM136444.

